# Evaluation of protein kinase D auto-phosphorylation as biomarker for NLRP3 inflammasome activation

**DOI:** 10.1101/2021.03.04.433858

**Authors:** Diane Heiser, Joëlle Rubert, Adeline Unterreiner, Claudine Maurer, Marion Kamke, Ursula Bodendorf, Christopher J. Farady, Ben Roediger, Frédéric Bornancin

**Author notes:** Corresponding author (FB).

## Abstract

**Background:** The NLRP3 inflammasome is a critical component of sterile inflammation, which is involved in many diseases. However, there is currently no known proximal biomarker for measuring NLRP3 activation in pathological conditions. Protein kinase D (PKD) has emerged as an important NLRP3 kinase that catalyzes the release of a phosphorylated NLRP3 species that is competent for inflammasome complex assembly.

**Methods:** To explore the potential for PKD activation to serve as a selective biomarker of the NLRP3 pathway, we tested various stimulatory conditions in THP-1 and U937 cell lines, probing the inflammasome space beyond NLRP3. We analyzed the correlation between PKD activation (monitored by its auto-phosphorylation) and functional inflammasome readouts.

**Results:** PKD activation/auto-phosphorylation always preceded cleavage of caspase-1 and gasdermin D, and treatment with the PKD inhibitor CRT0066101 could block NLRP3 inflammasome assembly and interleukin-1β production. Conversely, blocking NLRP3 either genetically or using the MCC950 inhibitor prevented PKD auto-phosphorylation, indicating a bidirectional functional crosstalk between NLRP3 and PKD. Further assessments of the pyrin and NLRC4 pathways, however, revealed that PKD auto-phosphorylation can be triggered by a broad range of stimuli unrelated to NLRP3 inflammasome assembly.

**Conclusion:** Although PKD and NLRP3 become functionally interconnected during NLRP3 activation, the promiscuous reactivity of PKD challenges its potential use for tracing the NLRP3 inflammasome pathway.

## Introduction

The NOD-like receptor (NLR) family, pyrin domain-containing protein 3 (NLRP3) inflammasome is involved in several pathophysiological conditions featuring sterile inflammation (1). Beyond a well-established transcriptional priming step for upregulation and expression of inflammasome components, several post-translational licensing events have been reported that allow for and contribute to NLRP3 inflammasome activation (2–5). Each inflammasome component, i.e., NLRP3, apoptosis-associated speck-like protein containing a CARD (ASC), Caspase-1, and NIMA-related kinase 7 (NEK7) can undergo post-translational modifications, such as phosphorylation, ubiquitination, acetylation or neddylation, but how these events are coordinated for effective inflammasome licensing and activation remains largely unknown.

Phosphorylation of NLRP3 has emerged as an important regulator for both functional maturation and acute activation of NLRP3 (6,7). Phosphorylation at serine residues in particular, was reported at three different sites of the human NLRP3 protein. Phosphorylation of S5 in the pyrin domain appears to be an early event. Removal of this phosphate group by protein phosphatase 2A (8) allows for subsequent activation steps. Phosphorylation of S198 between the pyrin and the NACHT domains, likely by c-Jun kinase (JNK)-1, is a critical licensing step (9). Phosphorylation/dephosphorylation at S295 in the NACHT domain appears to regulate membrane shuttling of NLRP3. Protein kinase A-mediated phosphorylation of this site is an early marker for cytoplasmic inactive NLRP3. Dephosphorylation is then required for activation at the membrane and subsequent re-phosphorylation by protein kinase D (PKD) allows for cytoplasmic release and association with ASC (10,11). Mutations at S295 were shown to cause a phenotype similar to that of activating NLRP3 mutations (cryopyrin-associated periodic syndrome), further illustrating the importance of this regulatory site (12).

Robust NLRP3 activation often requires transcriptional priming, in which mRNA and protein levels of inflammasome components, including those of NLRP3, are upregulated. Indeed, many studies have used increased levels of NLRP3 as supportive evidence for NLRP3 involvement. However, NLRP3 levels alone do not indicate activation status and more informative biomarkers are required to better assess the pathophysiological relevance of NLRP3 in diseased tissues. A critical step for activation of the NLRP3 inflammasome is its oligomerization followed by complex formation with ASC, Caspase-1 and NEK7. Polymerization of ASC is a hallmark of inflammasome activation and can be readily monitored (13–15). However, ASC polymerization itself does not inform about which specific inflammasome sensor molecule was involved at the assembly step. Monitoring NLRP3 phosphorylation for following NLRP3 maturation could potentially be an alternative. However, high quality anti-phospho NLRP3 antibodies are not presently available.

PKD has emerged as an important NLRP3 kinase that catalyzes the release of a phosphorylated NLRP3 species that is competent for inflammasome complex assembly (11). Here, we studied the functional relationship between PKD and NLRP3 with a biomarker perspective. We investigated the activation of PKD under various stimulatory conditions not limited to NLRP3, to probe the inflammasome space more broadly. Our data identified a strong functional interconnection between PKD and NLRP3 but also showed promiscuous reactivity of PKD, which challenges its potential use to sense NLRP3 functionality.

## Materials and Methods

### Chemicals

Nigericin (Enzo, BML-CA421-005), ouabain (Sigma #O3125), gramicidin (Sigma #G5002), niclosamide (Adipogen #AG-CR1-3643), DMSO, used as vehicle for compound dilutions (Sigma #D2650), MCC950 (either from Sigma, #5381200001 or synthesized at Novartis), CGP084892 and AFN700 (both synthesized at Novartis), CRT0066101 (Tocris #4975). The pyrin activating compound BAA473 has been previously described (16); it was synthesized at Novartis. The NLRC4 activating recombinant Salmonella Typhimurium Protein Prgl was from BioSource #MBS1177087.

### Antibodies

Anti-phospho PKD #2051, anti-PKD3 (D57E6) #5655, anti-PKD2 (D1A7) #8188, anti-human caspase-1 (D7F10) #3866, anti-cleaved Gasdermin-D (E7H9G) #36425, were from Cell Signaling Technologies. The anti-tubulin antibody (#T6074) was from Sigma. The anti-ASC antibody coupled to PE (#653904) used for ASC speck analysis by flow cytometry was from Biolegend.

### THP-1 cell lines and treatments

THP-1 cells (ATCC, TIB-202) were grown at 37°C 5% CO_2_ in medium (RPMI-1640 GlutaMAX^™^ + 25mM Hepes (Gibco #72400); 10% FBS (Gibco #16140); 1 mM Sodium Pyruvate (Gibco #11360); 0.05 mM 2-mercaptoethanol (Gibco#31350); 1x Pen/Strep (Gibco #15140); 100 µg/mL normocin (InvivoGen# ant-nr-1). Thirty µL of 5 x10E5 THP-1 cells/ml were seeded per well of a 384-well plate and incubated at 37°C 5%CO_2_ for 24h. Compounds were added (0.1% DMSO final concentration) and after one hour, cells were stimulated and incubated as specified in the figure legends. The NLRP3 KO THP-1 cell line, expressing an N-terminally truncated, inactive form of NLRP3, was from InvivoGen (thp-konlrp3z). The NLRC4 overexpressing THP-1 NLRC4 cells (InvivoGen, thp-nlrc4) were grown in THP-1 complete medium supplemented with 10 μg/ml blasticidin.

For NLRP3 inflammasome activation experiments measuring IL-1β production, priming with 100 nM PMA for 3h followed by overnight incubation without PMA was implemented, followed by activation with nigericin (15 µM unless stated otherwise) or other stimuli, for various time points specified in the figures. IL-1β levels in the supernatant were measured by HTRF (#62HIL1BPEH). Experiments reading out for ASC specks, Caspase-1 activation, and GSDMD cleavage were performed without the PMA priming step. For NLRC4 inflammasome activation, THP-1 cells and THP-1 NLRC4 cells were treated with 3 μg/ml of PrgI. For pyrin activation experiments, cells were treated with 100 μM BAA473.

### U937 cell lines and treatments

U937 cells were from ATCC (#CRL-1593). The pyrin-overexpressing U937 cell line has been previously described (16). Treatments were performed in a similar way to THP-1 cells, according to the details provided in the Figure legend.

### Flow cytometry

THP-1 cells were pretreated with 20 μM CGP084892, an irreversible caspase 1 inhibitor, to prevent nigericin-induced pyroptosis and subsequent ASC speck loss. After nigericin stimulation, THP-1 cells were treated for 30 min at 4°C in fixation buffer (4% formaldehyde (Sigma #47608)/5 mM EDTA) and then washed in permeabilization buffer (PBS without Ca^2+^/Mg^2+^ supplemented with 0.5% Triton-X100 and 5% FCS). The anti-ASC PE antibody was diluted in permeabilization buffer and added to cells at 1% final concentration. After an overnight incubation at 4°C and washing steps in permeabilization buffer, cells were finally resuspended in PBS and ASC specks were measured by FACS using a Canto II device, as described (15).

### Immunoblotting

THP-1 cells were collected in a 15 ml conical tube, spun down at 335 g for 5 min at 4°C. The cell pellet was washed with cold PBS and was resuspended afterwards in 1 vol. of RIPA buffer (#R0278, Sigma) supplemented with phosphatase inhibitors (Sigma, #P0044, #P5726) and cOmplete, EDTA-free protease inhibitor cocktail tablets (Roche, #11873580001). An equal volume of NuPage™ LDS sample buffer 4X (#NP0007, ThermoFisher) supplemented with 25% reducing agent (#NP0009, ThermoFisher) was mixed with the lysed cells suspension and denatured for 10 min at 95°C. Twenty µl samples and 4 µl PageRuler^™^ Plus ladder (#26619, ThermoFisher) were loaded on NuPage Bis Tris 4-12% 1.5 mm x 10 wells gels (ThermoFisher), migrated in MES SDS Running buffer (#NP0002, ThemoFisher) for 1h at 160V on ice. Proteins were transferred to PVDF membranes (#IB24001, ThermoFisher) on an iBlot^™^ device, program #3 for 7 min (ThermoFisher). The membrane was saturated by incubation in PBS (without EDTA) supplemented with 0.1% Tween 20 (#161-0781, BioRad) and 5% skimmed milk (Sigma, #70166). Primary antibodies were diluted 1:1000 in this buffer and incubated overnight at 4°C with gentle shaking. After washing steps using the same buffer, the secondary antibody (anti-Rabbit HRP CST #7074 or anti-Mouse HRP CST #7076) was diluted to 1:5000 and incubated for 1 hour at room temperature in PBS/Tween without milk. After final washes in this buffer, the membrane was incubated in ECL Immobilon^R^ Western (#WBKLS0500, Millipore) for 1 min and scanned using an ECL Fusion Fx imaging device.

## Results

### PKD auto-phosphorylation coincides with NLRP3 activation

NLRP3 is a sensor for cellular stress. Thus, a number of unrelated drugs known to cause cellular damage, ranging from bacterial toxins to crystals or biochemicals with defined mechanisms of action, are able to trigger the NLRP3 inflammasome (17–19). Among those, we selected four agents, based on their reported differences in strength and mechanism, namely nigericin, gramicidin, ouabain and niclosamide. Nigericin and gramicidin are microbial pore-forming toxins that activate the NLRP3 inflammasome due to their capacity to induce potassium efflux, a key step for triggering NLRP3 (20) ―Nigericin is a particularly strong activator because it induces additional changes such as mitochondrial perturbations (20). Ouabain is an inhibitor of the Na^+^-K^+^ ATPase that also leads to potassium efflux and NLRP3 activation. The proton uncoupling agent niclosamide more modestly stimulates NLRP3 via intracellular acidification and mitochondrial inhibition (21). We first compared these four stimuli by evaluating side-by-side their impact on IL-1β secretion, which is produced and released downstream of NLRP3 activation after cleavage of pro-IL-1β. All treatments resulted in IL-1β production but varied with respect to their kinetics and extent of IL-1β release. Nigericin showed a rapid onset rapidly reaching maximum levels of IL-1β production, consistent with its broad-acting pharmacology (Fig. 1A). By comparison, the onset was delayed for ouabain and gramicidin, leading to respectively high and medium levels of IL-1β production, while niclosamide was the least efficacious agent under our experimental conditions (Fig. 1A).

**Fig. 1:**
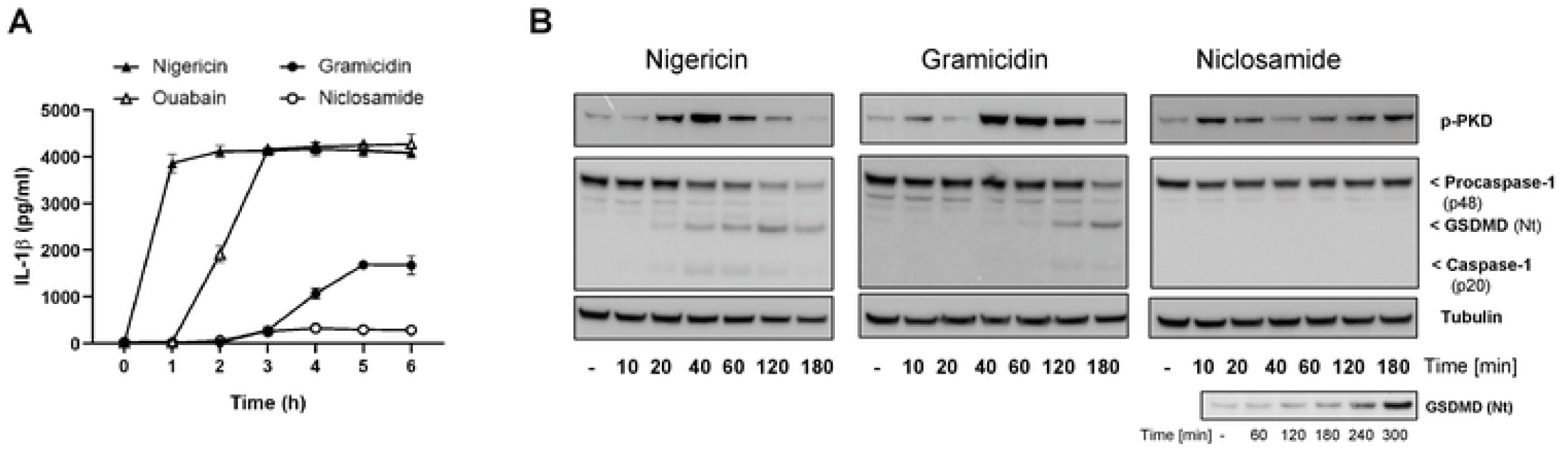
PKD auto-phosphorylation coincides with NLRP3 activation. (A) IL-1β secretion measured using THP-1 cells primed with 100 nM PMA and stimulated with nigericin (15 µM), ouabain (1 uM), gramicidin (10 uM) or niclosamide (1uM) after the indicated time course. This is one of thwo experiments with similar results. (B) Stimulation time course using THP-1 cells treated with either nigericin, gramicidin, or niclosamide followed by whole cell lysates analysis by immunoblotting to detect PKD activation (auto-phosphorylation) and NLRP3 inflammasome activation (caspase-1 and gasdermin D cleavage). Tubulin levels were used as loading controls. This is one of two (gramicidin) or three (nigericin, niclosamide) experiments with similar results. An additional experiment with a prolonged time course in the presence of nigericin was implemented to allow for detecting NLRP3 activation (lower panel).

PKD and its kinase activity are required for NLRP3 activation after stimulation by nigericin. This was previously established based on PKD knock-down and pharmacological approaches in bone marrow-derived macrophages and in THP-1 cells (11). We wondered whether differences in kinetics and efficiency of NLRP3 activators might have an impact on PKD activity. To test this, we looked at proximal NLRP3 activation readouts, i.e. caspase-1 and gasdermin D cleavage, and in parallel monitored PKD activity by following its auto-phosphorylation at S916 (22). Nigericin induced a time-dependent increase in cleaved caspase-1 (p20) and cleaved gasdermin D (GSDMD Nt) with a transient maximum at 40 and 120 min, respectively, followed by a decrease due to pyroptotic cell death (Figure 1B). Caspase-1 p20 and GSDMD Nt were also produced upon treatment with gramicidin, albeit with a slower onset as compared to nigericin (Figure 1B), consistent with the IL-1β secretion data (Figure 1A). Niclosamide treatment led to only limited GSDMD Nt levels, detectable only after prolonged kinetics and enhanced immunoblotting sensitivity (Figure 1B). Of note, auto-phosphorylation of PKD induced by nigericin or gramicidin coincided with NLRP3 activation, peaking closely to the time when GSDMD Nt started to be produced. Treatment with niclosamide consistently led to a biphasic pattern of PKD auto-phosphorylation. A first wave was detected that resoled within 30 minutes and was not linked to detectable NLRP3 activity. However, a second wave of PKD auto-phosphorylation started after 1h, continued until at least 3h, and coincided with the appearance of cleaved gasdermin D (Figure 1B).

These results suggested that auto-phosphorylation of PKD induced by NLRP3 inflammasome activators follows a pattern that reflects both the strength and the mechanism of the activator.

### PKD kinase inhibition down-regulates NLRP3 activation

To evaluate the contribution of PKD kinase activity to NLRP3 inflammasome activation, we used the pyrazine benzamide compound CRT0066101, a potent PKD inhibitor (23), which fully blocked PKD auto-phosphorylation induced by nigericin (Figure 2A). CRT0066101 was tested in parallel to MCC950, a compound known to prevent assembly of the NLRP3 inflammasome and thereby abolish IL-1β production after stimulation (24,25). A time course with 15 µM nigericin showed a strong dependency of NLRP3-induced ASC speck formation and IL-1β release on PKD activity at early time points (Fig. 2B). With increasing exposure times, the levels of pathway stimulation continued to rise while the overall outcome became less PKD-dependent, and at 3h blocking PKD no longer had an impact.

**Fig. 2:**
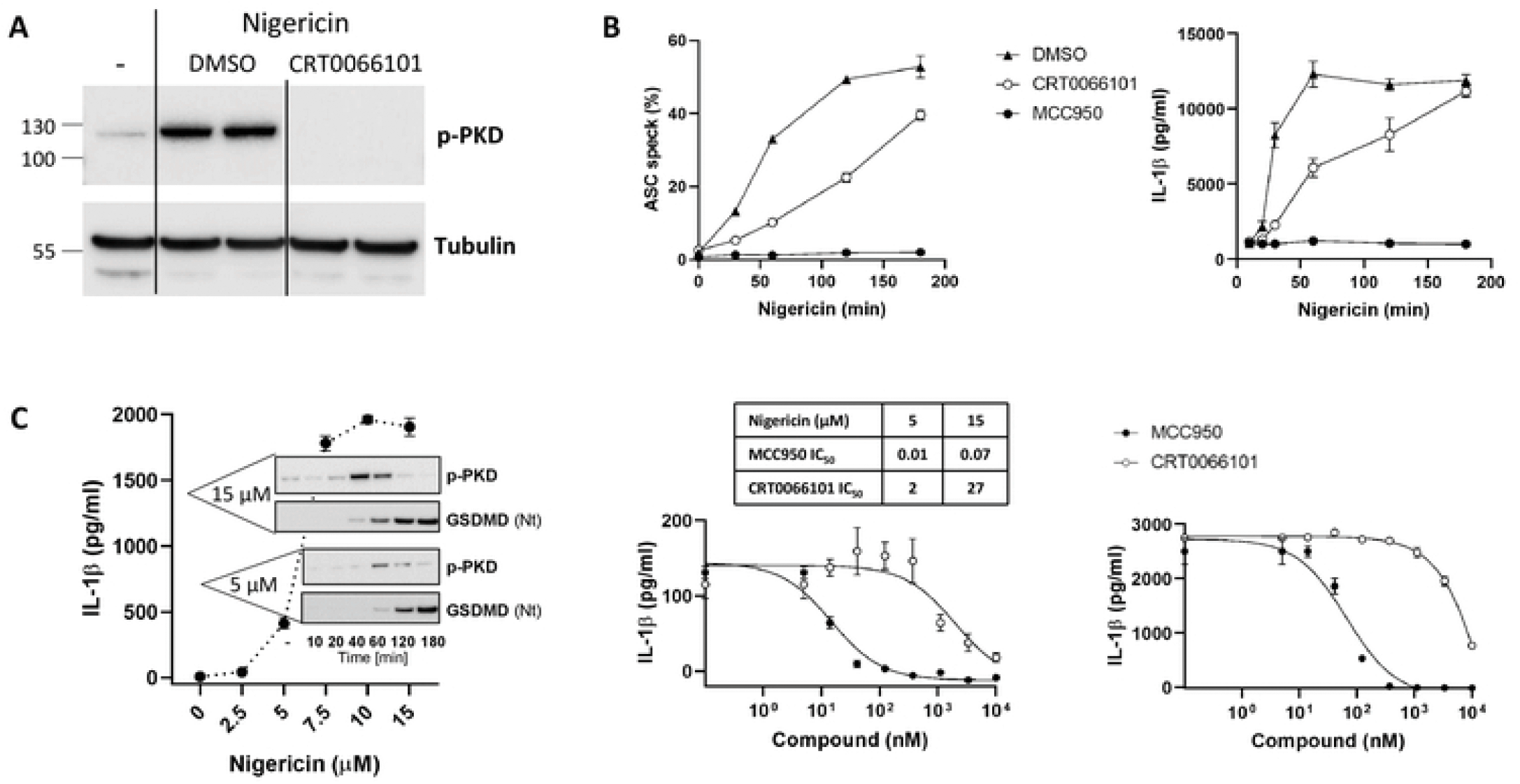
PKD kinase inhibition down-regulates NLRP3 activation. (A) THP-1 cells were treated with CRT0066101 (10 µM) or 0.1 % DMSO (vehicle control) for one hour before challenge with nigericin (15 µM) for 20 min. Immunoblot of whole cell lysates probed for auto-phosphorylated PKD. Tubulin is used as loading control. One of three experiments with similar results is shown (B) *Left*, FACS analysis of ASC speck formation in THP-1 cells treated with 15 uM nigericin for up to 3h in the presence of 20 µM CGP084892 (used to prevent pyroptosis and the resulting speck leakage), in the presence or absence of CRT0066101 (10 µM), MCC950 (1 µM) or vehicle control. Average values of two determinations ±SD. *Right*, IL-1β secretion measured using PMA-primed THP-1 cells, treated with CRT0066101 (10 µM), MCC950 (1 µM) or vehicle control for one hour, before a time course of stimulation with nigericin (15 µM). This is one of three experiments with similar results. (C) *Left*, IL-1β secretion measured in THP-1 cells after 3h of incubation with a concentration range of nigericin; corresponding immunoblots displaying auto-phosphorylated PKD and cleaved gasdermin D levels obtained upon a kinetics of stimulation with 5 µM and 15 µM nigericin. One of two similar experiments is shown. *Right*, Effect of concentration ranges of MCC950 and CRT0066101 on IL-1β secretion, tested upon stimulation with nigericin (5 µM for 5h, or 15 µM for 3h). Means of triplicate determinations ±SEM. IC_50_ (µM) for both compounds under both experimental conditions are indicated in the table.

To better understand the correlation between the dependency on PKD versus the strength of the activation stimulus, we selected a lower concentration of nigericin (5 uM), which triggered lower IL-1β release, following a slower onset and reduced PKD activation (Fig. 2C). Under these conditions, CRT0066101 blocked IL-1β release with an IC_50_ of 2 µM, compared to 27 µM when measured after stimulation with 15 µM nigericin (Fig. 2D). This shift towards a higher IC_50_ value was also observed with the NLRP3 inhibitor (14 nM after 5 µM nigericin, vs. 66 nM after 15 µM nigericin), which suggested that the potency of CRT0066101 might have been too low for continuous blockade under strong and prolonged NLRP3 activation. Given the ability of NLRP3 to act as a seed for polymerizing ASC, which leads to an amplification mechanism, any residual NLRP3 activation likely has profound consequences.

### NLRP3 blockade prevents PKD auto-phosphorylation

Phosphorylation of NLRP3 by PKD is one of the final regulatory steps during NLRP3 maturation, enabling NLRP3 to interact with ASC. Indeed, the impact of CRT0066101 on IL-1β after nigericin treatment was mirrored by its effect on ASC speck formation (Fig. 2B). Given the close spatio-temporal coordination between PKD activation and NLRP3 functional assembly, we asked whether regulation of PKD might be sensitive to NLRP3 functionality, that is, whether direct inhibition of NLRP3 might have an impact on PKD activity. We tested three principles, interfering at different stages of the NLRP3 assembly process. MCC950, which prevents oligomerization of NLRP3, efficiently blocked nigericin-induced PKD auto-phosphorylation (Fig. 3A). AFN700, an IKK inhibitor recently shown to blunt the efficiency of the inflammasome activation cascade through stabilization of procaspase-1 (Unterreiner et al., *accepted manuscript*), led to reduced PKD auto-phosphorylation (Fig.3A). Finally, CGP044892, a caspase-1 inhibitor (26) that is as effective as MCC950 in blocking the NLRP3 inflammasome, completely prevented PKD auto-phosphorylation (Fig. 3A). Therefore, on the one hand, PKD activation is required for NLRP3 function, and on the other hand, as shown collectively by these data, the ability of NLRP3 to assemble into a functional inflammasome appears to be key for sustaining PKD auto-phosphorylation.

**Fig. 3:**
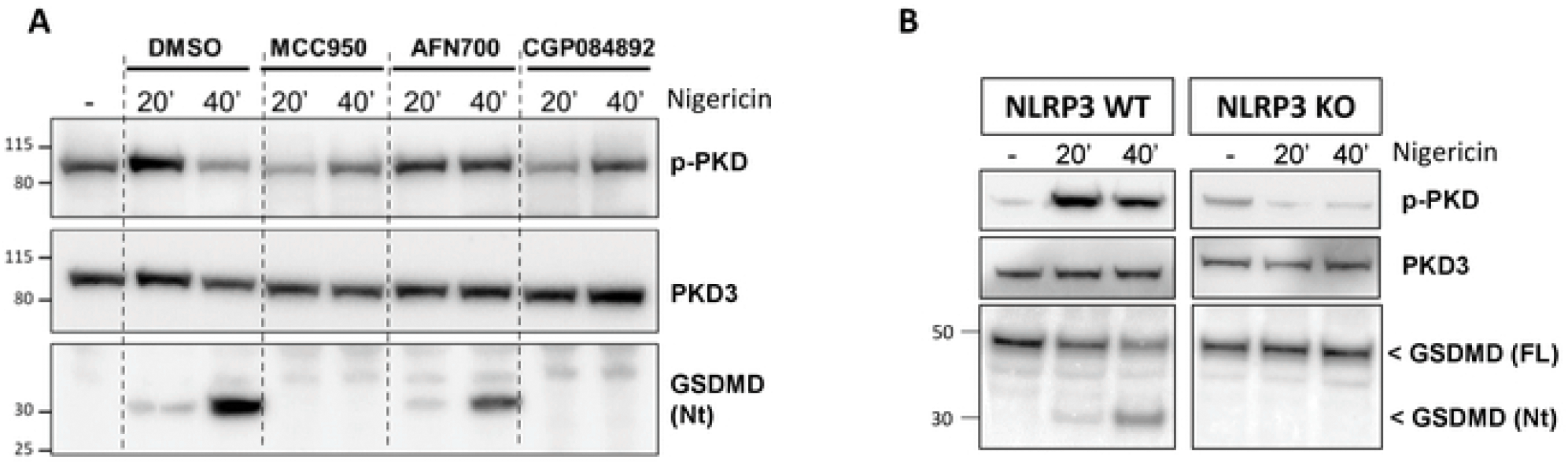
NLRP3 blockade prevents PKD auto-phosphorylation. (A) Immunoblot analysis of whole cell lysates from THP-1 cells treated with nigericin for the indicated times ± MCC950 (1 µM), AFN700 (3 µM) or CGP084892 (20 µM), monitoring total and auto-phosphorylated PKD, as well as gasdermin D cleavage. THP-1 were reported to selectively express PKD3 (31). (B) Immunoblot analysis of whole cell lysates from THP-1-WT or NLRP3-KO cells treated with nigericin for the indicated times, monitoring total and auto-phosphorylated PKD, as well as full and N-terminally cleaved gasdermin D.

To obtain a more direct evidence of a functional crosstalk between PKD and NLRP3, we used a THP-1 cell line expressing a non-functional N-terminally truncated variant form of NLRP3. Nigericin did not induce cleavage of GSDMD in this line and, remarkably, it was unable to raise the levels of auto-phosphorylated PKD, in contrast with its effect in the wild-type line (Fig.3B).

### PKD auto-phosphorylation is not a selective inflammasome marker

The functional relationship identified between PKD and NLRP3 made us wonder whether this was a specific feature of this particular inflammasome. To address this question, we extended our investigations to two additional inflammasomes, the NLR family, CARD domain-containing protein 4 (NLRC4), and pyrin. We first used control and NLRC4-ovexpressing THP-1 cells and stimulated them with *Salmonella typhimurium* PrgI protein. GSDMD cleavage was detectable after as little as 5 min treatment, in both WT and NLRC4-overexpressing cells, with a stronger signal in the latter (Fig. 4A). In both lines, the appearance of cleaved GSDMD coincided with higher levels of auto-phosphorylated PKD (Fig. 4A). To rule out a possible cooperation of NLRP3, we treated THP-1 cells with MCC950 prior to NLRC4 activation. MCC950 did not prevent GSDMD cleavage and actually increased PKD auto-phosphorylation levels, suggesting that the NLRC4 activator had an impact on PKD auto-phosphorylation, independently of NLRP3 (Fig.4B). The same rationale was applied to probe selectivity versus the pyrin inflammasome. In this case we stimulated THP-1 cells with the bile acid analogue BAA473 (16), and we tested the effect of MCC950. Similarly to what was observed for NLRC4 activation, the pyrin inflammasome activator BAA473 induced GSDMD cleavage shortly following on PKD auto-phosphorylation, and here as well, MCC950 was not inhibitory (Figure 4C). These findings were reproduced in U937 monocytic overexpressing pyrin (Figure 4D). These experiments thus indicated that PKD auto-phosphorylation can occur upon pan-inflammasome stimulation implying it is not a selective marker for NLRP3 pathway engagement.

**Fig. 4:**
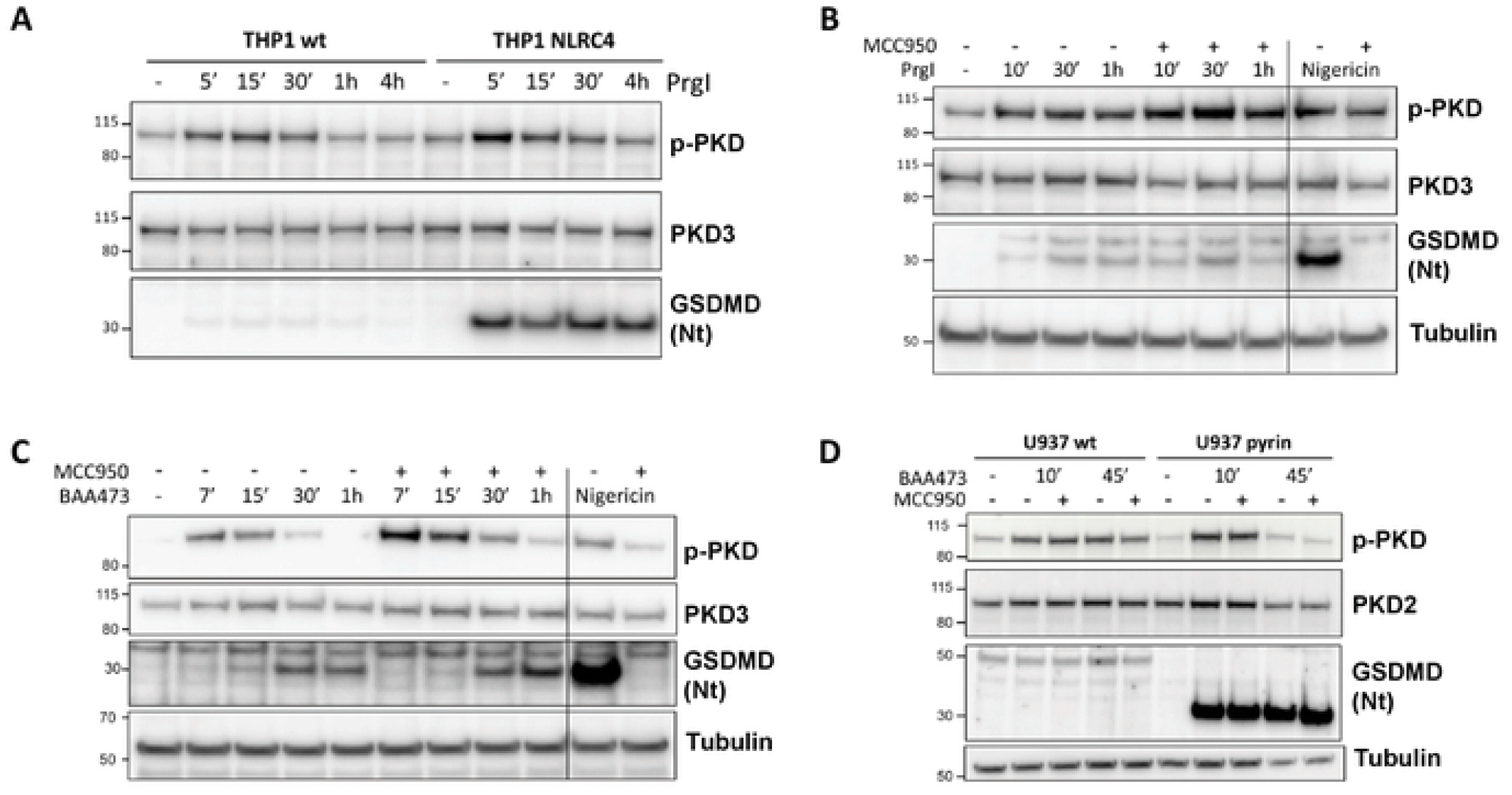
PKD auto-phosphorylation is not a selective inflammasome marker. (A) Time course of stimulation of THP1-WT and THP1-NLRC4 cells upon stimulation with PrgI (3 µg/ml). Immunoblot analysis of whole cell lysates showing total and auto-phosphorylated PKD, as well as gasdermin D cleavage. (B) Time course of stimulation of THP1-WT with PrgI (3 µg/ml) ± MCC950 (10 µM). Nigericin stimulation ± MCC950 (10 µM) serves as positive control. Immunoblot analysis of whole cell lysates showing total and auto-phosphorylated PKD, as well as gasdermin D cleavage. Tubulin levels are used for comparison of PrgI and nigericin stimulatory conditions. (C) Time course of stimulation of THP1-WT with BAA473 (100 µM) ± MCC950 (10 µM). Nigericin stimulation ± MCC950 (10 µM) serves as postive control. Immunoblot analysis of whole cell lysates showing total and auto-phosphorylated PKD, as well as gasdermin D cleavage. Tubulin levels are used for comparison of BAA473 and nigericin stimulatory conditions. (D) Time course of stimulation of U937-WT and U937-Pyrin cells with BAA473 (100 µM) ± MCC950 (10 µM). Immunoblot analysis of whole cell lysates showing total and auto-phosphorylated PKD, as well as gasdermin D cleavage. PKD2 levels were monitored as PKD1/3 were not detectable in U937 cells. Lower tubulin levels in pyrin-overexpressing cells at 45 min account for cell death triggered by BAA473.

## Discussion

PKD is a family of enzymes responding to diacylglycerol, which is produced downstream of several receptor tyrosine kinases and G-protein coupled receptors (27). Thus, regulation of PKD activity can clearly take place independently of inflammasome signaling. Our interest for PKD in the context of the NLRP3 inflammasome was based on three parameters. First, the reported requirement for PKD to activate NLRP3 (11). Second, the importance of PKD enzymes for trafficking from the trans-Golgi network, a cellular compartment, which, upon treatment with NLRP3 activators becomes closely associated with mitochondria-associated endoplasmic reticulum membranes where NLRP3 is located (11). Third, the recent evidence for PKD dimerization and activation, which may take place at various subcellular locations including the trans-Golgi network (28), where diacylglycerol is known to be produced upon NLRP3 activation (11).

NLRP3 becomes competent for inflammasome formation after a multi-step maturation process involving post-translational modifications. In this work, we showed that PKD, a kinase required for the final permissive phosphorylation of NLRP3 at serine 295, is controlled by NLRP3 itself, suggesting that a functional crosstalk between PKD and NLRP3 may operate. Given that no antibody tool is currently available to trace p-295 NLRP3, and because evaluation of NLRP3 activity in biological samples has remained challenging, this prompted us to consider whether information on PKD activation levels might be used as a surrogate for NLRP3 activation.

PKD activation also occurred upon stimulation of other inflammasomes, such as NLRC4 and pyrin, and we showed this was not a consequence of possible co-opting of NLRP3, which in known to occur under some circumstances (29). Pyrin is known to traffic along microtubules to reach the centrosome region using a similar mechanism as NLRP3 (30). However, NLRC4 biology is more divergent. Therefore, the stimulation of PKD obtained under broad inflammasome activation conditions likely results from promiscuous reactivity of PKD to cellular challenge.

## Conclusions

The present data do not support monitoring PKD auto-phosphorylation as a means to identify NLRP3 activation even though they do not exclude the possibility that specific subsets of PKD may selectively engage with NLRP3 during activation.

## Abbreviations

CARD: (caspase activation and recruitment domain)
ASC: (apoptosis-associated speck-like protein containing a CARD)
NOD: (nucleotide-binding oligomerization domain)
NACHT: [(NAIP (neuronal apoptosis inhibitory protein)
CIITA: (MHC class II transcription activator)
HET-E: (incompatibility locus protein from Podospora anserina) and
TP1: (telomerase-associated protein)]
NLRP3: (NOD-like receptor family
pyrin: domain-containing protein 3)
NLRC4: (NLR family
CARD: domain-containing protein 4)
PKD: (protein kinase D)
c-Jun kinase: (JNK)
Gasdermin D: (GSDMD).

## Declarations

### Data Availability Statement

All relevant data are within the paper.

## Acknowledgements

We thank Irina Alimov for provision of the U937-Pyrin expressing cell line, and Tim Schuhmann and Anne Schitter for synthesis of the bile acid analogue BAA473.

## Author Contributions

**Conceptualization:** Frederic Bornancin.

**Data curation:** Diane Heiser, Joëlle Rubert, Adeline Unterreiner.

**Formal analysis:** Diane Heiser, Joëlle Rubert, Adeline Unterreiner, Ursula Bodendorf, Christopher J Farady, Ben Roediger, Frederic Bornancin.

**Investigation:** Diane Heiser, Joëlle Rubert, Adeline Unterreiner, Frederic Bornancin.

**Methodology:** Diane Heiser, Joëlle Rubert, Adeline Unterreiner, Claudine Maurer, Marion Kamke.

**Supervision:** Frederic Bornancin.

**Writing – original draft:** Frederic Bornancin.

**Writing – review & editing:** Ben Roediger, Frederic Bornancin.

